# Vγ usage distinguishes pro- and anti-tumor intestinal γδ T cell subsets

**DOI:** 10.1101/2021.11.13.468487

**Authors:** Bernardo S. Reis, Patrick W. Darcy, Iasha Z. Khan, Olawale Eleso, Caixia Zhu, Marina Schernthanner, Ainsley Lockhart, Aubrey Reed, Juliana Bortolatto, Tiago B. R. Castro, Angelina M. Bilate, Sergei Grivennikov, Daniel Mucida

## Abstract

γδ T cells physiologically scan the intestinal epithelium, representing a substantial fraction of infiltrating lymphocytes in colorectal cancer (CRC), albeit their role in CRC remains unclear. Using murine CRC models, we found that most γδ T cells in pre- or non-tumor colon express Vγ1^+^ or Vγ7^+^ and exhibit a cytotoxic profile. Targeting these γδ T cell subsets, as well as conditionally interfering with γδ T cell function at early stages of tumorigenesis led to heightened tumor development, suggesting anti-CRC functions for Vγ1^+^ and Vγ7^+^ subsets. In contrast, RORγt^+^ γδ T cell subsets, including Vγ4^+^ and microbiota-dependent Vγ6^+^, accumulated during CRC progression. Conditional deletion of RORγt or Vγ chains revealed redundant roles for IL-17–producing Vγ4^+^ and Vγ6^+^ γδ T cells in promoting tumor growth. Our results uncover pro- and anti-tumor roles for γδ T cell subsets.

---

Intestinal intraepithelial lymphocytes (IELs) comprise a large T cell population located at the critical interface between the core of the body and the intestinal lumen, which is constantly exposed to food, commensal microbes, and pathogens. Conversely, previous observations suggest an important role for IELs as a first line of immunity against pathogens in both mice and humans (*1–4*). Among the main IEL subsets in mice or humans, T cells harboring the γδ T cell receptor (TCR) display epithelial surveillance and antipathogen activities that are modulated through crosstalk with the intestinal epithelium, highlighting that γδ IELs are finely tuned to local epithelial signals (*3, 5*). In addition to their role in immune surveillance against enteric infections, sequencing analysis points to a γδ T cell gene signature as the most favorable prognostic factor among tumor-infiltrating leukocytes across cancer types, including colorectal cancer (CRC) (*6*). CRC is the second most deadly cancer in the United States, affecting over 140,000 people each year, killing approximately 50,000 in the US (ACS Inc., 2020). The cumulative risk of IBD patients to develop CRC can reach 20%, although most CRCs develop in patients without underlying inflammation. In both cases, tumor-elicited inflammation triggered by epithelial disturbances and microbial invasion is essential for survival of transformed cells, and tumor growth (*7–10*). Here, we addressed whether epithelial surveillance by γδ T cell subsets helps prevent damaged ECs from progressing into cancer and whether additional γδ T cell subsets play a contrasting role, accelerating tumor progression.

To assess the role of γδ T cells in CRC, we employed two distinct CRC-mouse models: the chemical AOM-DSS colitis-associated cancer (CAC) (*11*) and the genetically inducible APC deficiency (*9, 12, 13*) (Fig. 1A). Our initial characterization of γδ T cells in naïve animals confirmed (*14–16*) a dominance by Vγ1 and Vγ7-expressing subsets, followed by Vγ4^+^ and Vγ6^+^ subsets at a low frequency, and changes in the expression of CD8αα homodimers according to proximal-distal and villus-crypt axes (fig. S1A to C). In colitis-associated CRC (AOM-DSS), we observed that tumor-infiltrating γδ T cells adopt both anti- and pro-tumor phenotypes, with increased expression of CD107α and IFN-γ [associated with anti-tumor responses (*8, 17*)] as well as IL-17 [associated with pro-tumorigenic function (*9, 17*)](Fig. 1B). CD4^+^ TCRαβ^+^ cells also display IFN-γ and IL-17–producing phenotypes, but in contrast to γδ T cells, expression levels upon restimulation are consistent between tumor and non-tumor tissue sites (fig. S1D). In the CAC model, roughly 40% of IL-17–producing cells within the tumor are γδ T cells, while about 30% are CD4^+^ T cells (Th17 cells), while in non-tumor areas they correspond to roughly 35% and 20%, respectively (fig. S1E). Similar to what was observed in naïve mice, γδ T cells in healthy colon tissue display high expression of CD8αα, while a CD8α^-^ population expressing the exhaustion marker PD-1^+^ is prominent among tumor–infiltrating γδ T cells (Fig. 1C and 1D). PD-1 expression segregated cells with potential anti-tumor (PD-1^-^:IFN-γ^+^ and CD107^+^) or pro-tumor phenotype (PD-1^+^: IL-17^+^, IFN-γ^-^ and CD107^-^) (Fig. 1E and 1F). These findings were replicated in the inducible APC loss-based CRC model, where γδ T cells expressing IFN-γ and IL-17 also accumulate among tumor-infiltrating lymphocytes (Fig. 1G). As in the CAC model, tumor infiltrating PD-1^+^ (CD8α^-^) preferentially express IL-17, while PD-1^-^ (CD8a^+^) preferentially secrete IFN-γ (Fig. 1H to K). CD4^+^ T cells again do not show significant differences in IL-17 and IFN-γ production between tumor or adjacent non-tumor areas (fig S1F). In the APC loss model, about 60% of IL-17– producing cells within the tumor are γδ T cells, while Th17 cells constitute only 20%; outside tumor areas, almost 70% IL-17–producing cells are γδ T cells and roughly 10% are Th17 (fig. S1G). To further characterize tumor-infiltrating γδ T cells in the CAC model, we sorted these cells based on PD-1 expression and performed RNAseq analysis. PD-1^-^ γδ T cells (blue) display increased expression of cytotoxic related genes (e.g. *Gzmk, Crtam, Klra5*), while PD-1^+^ γδ T cells (red) show increased expression of genes associated with IL-17 and pro-tumorigenic responses (*10*) such as *Tmem176a* and *Il1r1* (Fig. 1L). These results point to opposing phenotypes between γδ T cells found at steady state or in non-tumor areas, and γδ T cells that accumulate during CRC.

**Figure 1.**
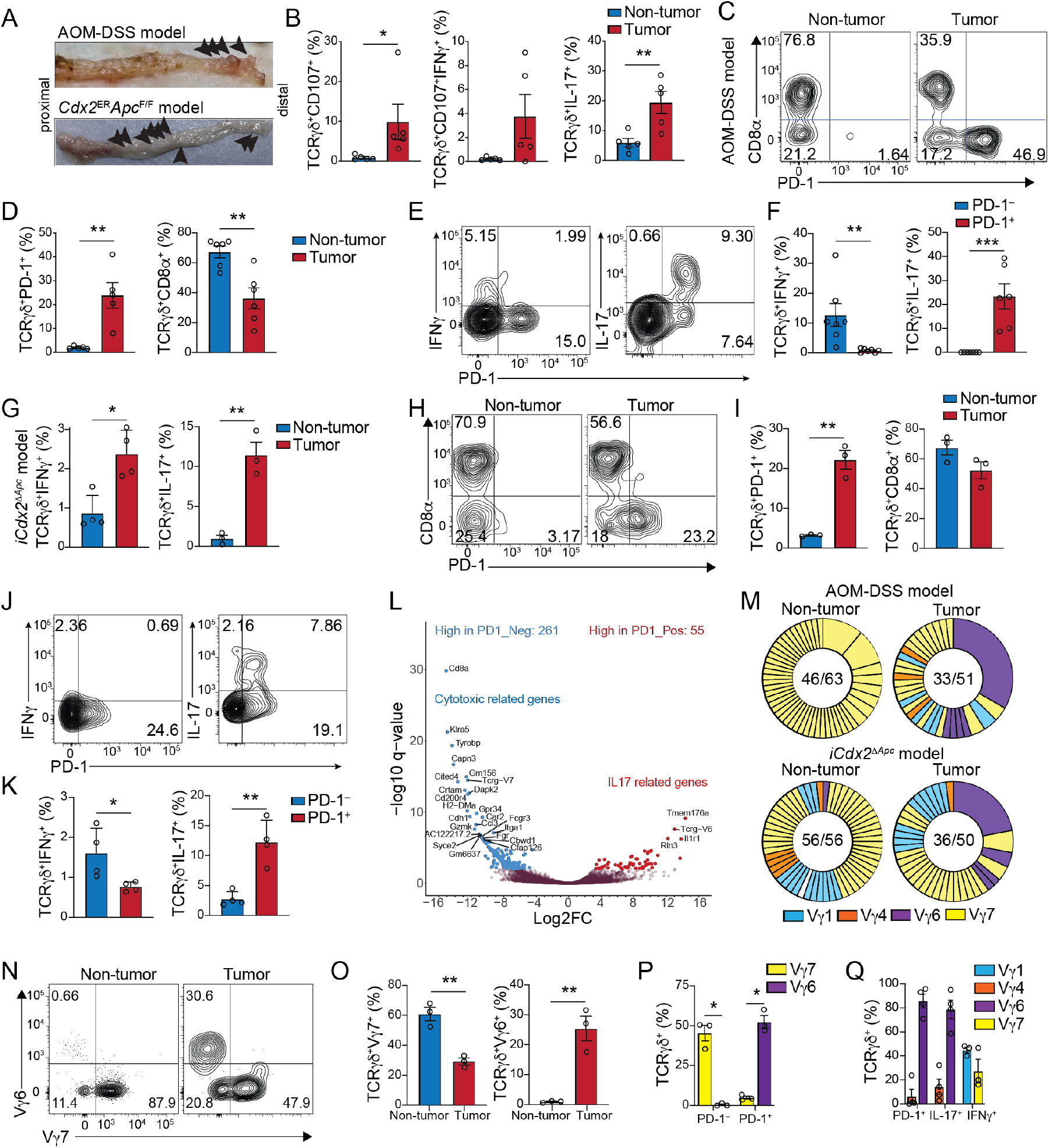
Profiling tumor-infiltrating γδ T cells in CRC models reveals distinct subsets. **(A)** Representative images of colon containing tumors (arrows) from wild-type mice injected with 12.5mg/kg of Azoxymethane (AOM) 2 days prior to 3 rounds of: 1 week “on” and 2 weeks “off’ 2% Dextran sodium sulfate (DSS) in the drinking water (top panel); or iCdx2^ΔAPC^ mice 5 weeks after 2 doses of 4mg/ml tamoxifen i.p. injection (lower panel). **(B-F)** Flow cytometry analysis of γδ T cells from tumor or non-tumor colonic tissue of mice subjected to the AOM-DSS model. **(B)** Frequency of CD107+, IFN-γ^+^CD107^+^ double positive, and IL-17^+^ among TCRγδ^+^ cells. **(C)** Representative dot-plot and **(D)** frequency of CD8α^+^ and PD-1^+^ among TCRγδ^+^ cells. **(E)** Representative dot-plot and **(F)** frequency of IFN-γ^+^ (left) or IL-17^+^ (right) among tumor-infiltrating PD-1^+^ or PD-F TCRγδ^+^ cells. **(G-K)** Flow cytometry analysis of γδ T cells from tumor or non-tumor colonic tissue of tamoxifen-treated iCdx2^ΔAPC^ mice. (G) Frequency of IFN-γ^+^ (left) and IL-17^+^ (right) among TCRγδ^+^ cells. (H) Representative dot-plot and **(I)** frequency of CD8α^+^ and PD-1^+^ among TCRγδ^+^ cells. **(J)** Representative dot-plot and **(K)** frequency of IFN-γ^+^ (left) or IL-17+ (right) among tumor-infiltrating PD-F or PD-F TCRγδ^+^ cells. (L) Volcano plot of differentially expressed genes from RNAseq analysis of sorted PD-F (blue) or PD-F (red) TCRγδ^+^ cells isolated from tumors of mice subjected to the AOM-DSS model. (M) Single-cell TCR sequencing of γδ T cells from tumor or non-tumor colonic tissue of mice subjected to the AOM-DSS (top) and APC loss (bottom) models. Numbers in the center of pie charts represent number of clones per total cells sequenced. Expanded clones are fused. Clones are colored based on Vγ usage. Purple clones represent expanded Vγ6Vδ4 cells. (N-Q) Vγ usage by γδ T cells from tumor or non-tumor colonic tissue of mice subjected to the AOM-DSS model. Representative dot-plot (N) and frequency (O) of Vγ6^+^ and Vγ7^+^ among TCRγδ^+^ cells. (P) Frequency of Vγ6^+^ and Vγ7^+^ among PD-F or PD-F TCRγδ^+^ cells from tumor-containing colonic tissue. (Q) Frequency of VγF, Vγ4^+^, Vγ6^+^ and Vγ7^+^ among tumor-infiltrating TCRγδ^+^ cells expressing PD-1, IL-17 or IFN-γ (AOM+DSS model). Representative data from 2 experiments with 3-4 animals per group. RNAseq and TCRseq data from pooled tumors. *P < 0.05). **P < 0.01, *** P<0.001. **(P, Q)** one-way ANOVA with Dunnett’s multiple comparison test); (B-O) two-tailed t-test. Error bars indicate SEM.

Our bulk RNAseq analysis suggested divergent TCR usage between tumor-infiltrating PD-1^+^ enriched in *TcrgV6* transcripts, and PD-1^-^γδ T cells, enriched in *TcrgV7* transcripts (Fig. 1L). To directly investigate the TCR repertoire of tumor–infiltrating γδ T cells, we single-cell sorted γδ T cells from tumor and adjacent non-tumor areas from mice subjected to AOM-DSS or APC loss CRC models. TCRseq analysis indicated that in the APC loss model, γδ T cells from non-tumor areas are completely diverse while tumor-infiltrating γδ T cells show noticeable clonal expansions (Fig. 1M). In the AOM-DSS model, some clonal expansion was seen in non-tumor areas likely due to a chronic inflammatory state (*18*), with slightly increased expansion observed in tumor-areas (Fig. 1M). Notably, the expanded clones found in tumor areas in both models were primarily Vγ6^+^Vδ4^+^clones rarely seen in non-tumor areas (Fig. 1M and fig. S1H and 1I). Flow cytometry analysis confirmed the relative increase of Vγ6^+^ cells at the expense of decreased Vγ7^+^ γδ T cells in tumor areas (Fig. 1N and 1O). As suggested by the bulk RNAseq analysis, tumor-infiltrating Vγ7^+^ T cells isolated from mice subjected to AOM-DSS are mostly PD-1^-^, while Vγ6^+^ cells express PD-1 (Fig. 1P). Overall, tumor-infiltrating PD-1^+^ and IL-17^+^ γδ T cells are composed of Vγ6 (85.7% ±5.6 and 78.8% ±6.4, respectively), and Vγ4 (6.5% ±5.7 and 14.2% ±6.4, respectively) while IFN-γ^+^ γδ T cells are composed of Vγ1 (44.9% ±2.4) and Vγ7 (27.5% ±9.6) in the CAC model (Fig. 1Q). Similar Vγ distribution was observed in tumor infiltrating TCRγδ^+^ T cells isolated from mice subjected to the APC loss model (fig. S1J). Vγ4^+^ and Vγ6^+^ subsets were previously described to leave the thymus from E15.5 and E18.5 pre-committed to IL-17 production, accumulating in specific sites such as the lung, genital tract, and adipose tissue (*19, 20*). The analyses above suggest that tumor infiltrating γδ T cells are composed of four major subsets based on Vγ-usage, divided into two functional groups: polyclonal Vγ7^+^ and Vγ1^+^ resembling colonic epitheliumresident γδ T cell subsets seen at steady state and exhibiting an “anti-tumor” cytotoxic program; and clonally-expanded Vγ6^+^Vδ4^+^ and at lower frequency, Vγ4^+^ γδ T cells, that express PD-1 and secrete IL-17.

To address whether the anti-tumor phenotype of epithelium-resident TCRγδ T cells plays a role in restricting CRC development, we first subjected *Trdc*^-/-^ knockout mice, which are deficient in γδ T cells, to AOM-DSS (Fig. 2A to C). Because *Trdc*^-/-^ animals are more susceptible to the DSS regimen (*21*), we used a lower DSS concentration (1%), which does not result in noticeable inflammation or tumor development in wild-type control mice (Fig. 2B and fig. S2A to C). Following the 1% DSS regimen, *Trdc*^-/-^ mice displayed enhanced inflammation and significant increase in tumor development, again suggesting an anti-tumor, or anti-inflammatory role for intestinal epithelium-resident γδ T cell populations (Fig. 2B and 2C). To extend these observations to a model which is not dependent on colitis, we lethally irradiated i*Cdx2*^ΔAPC^ recipient mice and transferred bone marrow cells from *Trdc*^-/-^ or wild-type control donor mice. Eight weeks after bone-marrow transplant, recipient mice were treated with tamoxifen to induce tumor development and analyzed 10 days later (fig. S2D). Consistent with the observations made in the AOM-DSS, i*Cdx2*^ΔAPC^ recipients of *Trdc*^-/-^ bone marrow show higher fecal lipocalin-2 and tumor incidence than mice recipient of wild-type bone marrow (fig. S2E and 2F). In these experiments, we opted to analyze tumor incidence instead of numbers given the difficulty in separating many joined tumors in mice recipient of *Trdc*^-/-^ bone marrow. Vγ6^+^ are canonical uterine γδ T cells known to leave thymus early during ontogeny, hence do not reconstitute in these bone-marrow transfer settings (*19, 22*). However, we observed reconstitution of CD8aa^+^ IFN-γ–producing populations (Vγ1^+^ and Vγ7^+^) and Vγ4^+^ γδ T cells in mice recipient of wild-type bone marrow (fig. S2G and S2H). The above data reinforce the possibility of an antitumor role for epithelial-resident γδ T cells.

**Figure 2.**
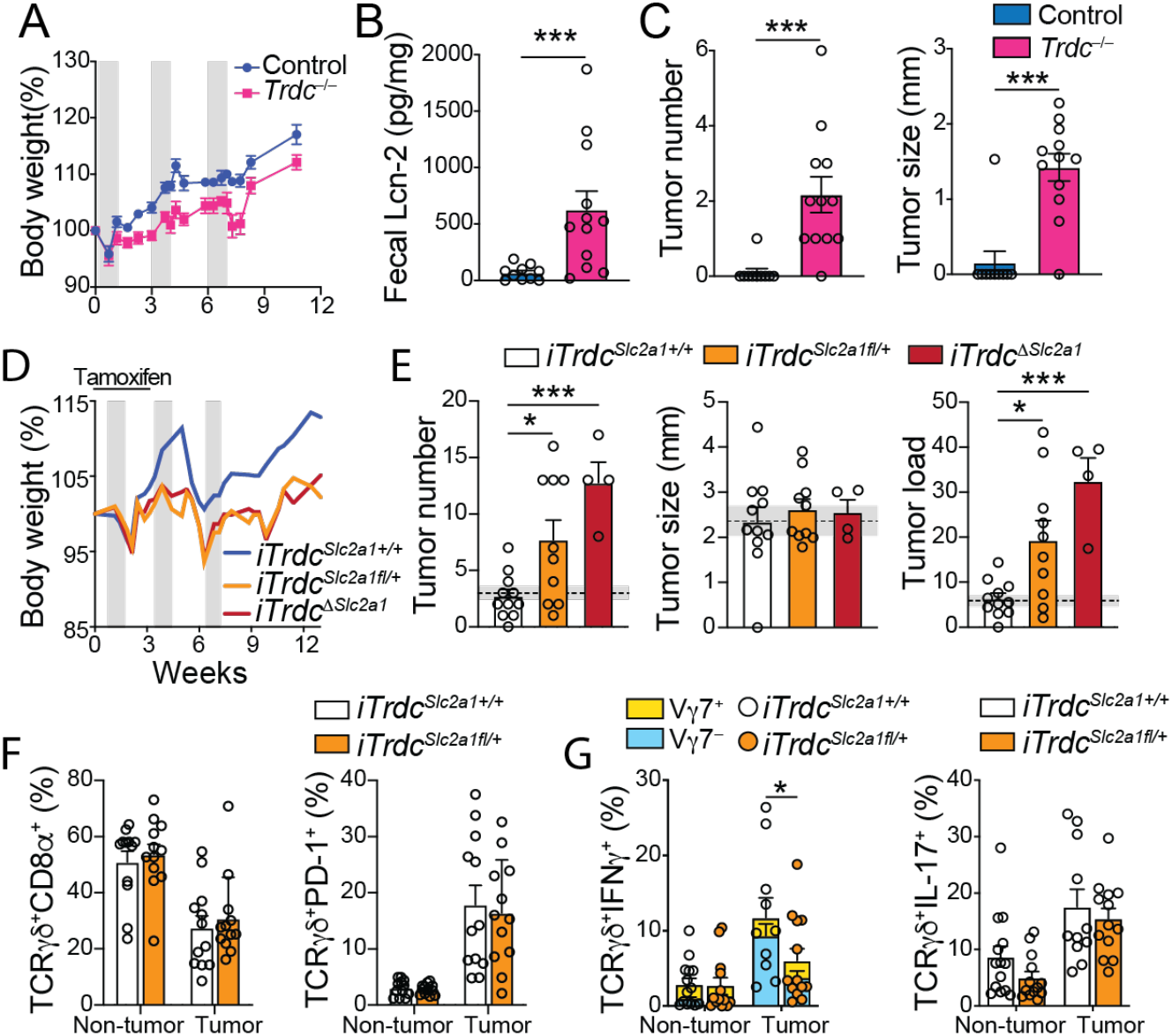
Loss-of-function by epithelium-resident γδ T cells results in increased tumor numbers. **(A-C)** *Trdc^-/-^* and littermate control mice were subjected to AOM-DSS model using low (1%) DSS dose and analyzed 12 weeks after initial AOM injection. **(A)** Percentage of body weight changes during AOM-DSS treatment. Gray bars represent DSS treatment. **(B)** Fecal lipocalin-2 quantification at week 12. **(C)** Tumor number and size. **(D-G)** i*TrdC*^Δ*Scl2a1*f/+^ or *iTrdC*^Δ*Scl2a1*^ and littermate control *iTrdc^Slc2a1+/+^*) mice were subjected to AOM-DSS model, treated with tamoxifen for the first 3 weeks and analyzed 12 weeks after initial AOM injection. *iTrdc^ΔSlc2a^* were only used for tumor quantification. **(D)** Percentage of body weight changes during AOM-DSS treatment. Gray bars represent DSS treatment. (E) Tumor number, size and load. Shaded area bounded by dashed lines indicates mean ± SEM of all control C57BL6/J mice analyzed in fig. S2B (AOM-DSS model). **(F-G)** Flow cytometry analysis of γδ T cells from tumor or non-tumor colonic tissue of i*TrdC*^Δ*Scl2a1*f/+^ and littermate control (i*TrdC*^Δ*Scl2a1*+/+^) mice subjected to AOM-DSS model. **(F)** Frequency of CD8oT (left) and PD-1^+^ (right), **(G)** IFN-γ^+^ (left) and IL-17^+^ (right) among TCRγδ^+^ cells in tumor or non-tumor colonic tissue. Vγ7^+^ (yellow) vs Vγ7-(blue) contribution to IFN-γ-producing γδ T cells is also shown. *Trdc^-/-^* and i*TrdC*^Δ*Scl2a1*f/+^ data are pooled from 3 experiments with 3-4 animals per group. *iTrdc^ΔSlc2a1^* data are from 1 experiment with 4 animals per group. *P < 0.05), **P < 0.01, ***P < 0.001. **(E-G)** One-way ANOVA with Dunnett’s multiple comparison test; **(B, C)** two-tailed t-test. Error bars indicate SEM.

Because total *Trdc*-deficiency also prevents the accumulation of γδ cells during tumor progression, we next aimed to preferentially restrict epithelium-resident γδ T cell function by inducible hemizygous or homozygous inactivation of *Scl2a1*[i*Trdc*^*Scl*2a1f+^ or *iTrdc^ΔScl2a1^*) encoding the glucose transporter Glut1. We have previously reported that epithelium-resident γδ T cells control early invasion by *Salmonella* Typhimurium via a metabolic switch towards glycolysis that is dependent on Glut1 expression (*3*), an observation related to recent findings showing that IFN-γ–, but not IL-17–secreting γδ T cells, are glycolytic and exert anti-tumor activity (*23*). We subjected i*TrdC*^Δ*Scl2a1*^, i*TrdC*^Δ*Scl2a1*f/+^, and littermate control mice to the AOM-DSS model (Fig. 2D). Early targeting of Glut1 in γδ T cells results in higher tumor number and load when compared to control animals, without affecting tumor size (Fig. 2E). Moreover, while Glut1 inactivation does not lead to changes in CD8α, PD-1 or IL-17 expression by γδ T cells, we detected a significant reduction in IFN-γ production (48.6%) by tumor-infiltrating γδ T cells in i*TrdC*^Δ*Scl2a1*f/+^ mice (Fig. 2F and 2G and fig. S2I). Within IFN-γ–producing γδ cells, Glut1 inactivation had a preferential impact on Vγ7^-^ cells (primarily represented by Vγ1^+^) (Fig. 2G). Among total tumor–infiltrating γδ T cells in tamoxifen-treated i*TrdC*^Δ*Scl2a1*f/+^ mice we find a significant reduction in Vγ7 (fig. S2J), overall indicating that inactivation of Glut1 in γδ T cells during early states of tumorigenesis mostly affects epithelium-resident Vγ1^+^ and Vγ7^+^ cells. In contrast, late Glut1 targeting (after the second DSS cycle) had no impact on tumor load or tumor-infiltrating γδ T cells (fig. S2K to O). These results suggest that epithelium-resident γδ T cells exert immunosurveillance against epithelial tumors and early impairment of Glut1–dependent γδ T cells favors CRC development.

To directly address the possible role of epithelium-resident γδ T cells in the regulation of tumor development, we generated Vγ7^-/-^ mice by CRISPR targeting of *Trgv7* gene. Fully backcrossed Vγ7^-/-^ mice were subjected to AOM-DSS and show no differences in tumor number, tumor size or colonic γδ T cells (aside from a reduction in Vγ7+ and CD8aa^+^ γδ T cells) when compared to heterozygous littermate controls (fig. S3A to E). Although, overall CD4^+^ T cells responses were unaffected, Vγ7^-/-^ mice displayed a reduction in IL-17–secreting CD4^+^ T cells in non-tumor areas when compared to Vγ7^+/-^ controls (fig. S3F). While the ratio of γδ/αβ T cells in non-tumor areas was significantly reduced, we observed near doubling of Vγ1^+^ cells in Vγ7^-/-^ mice when compared to Vγ7^+/-^ controls (fig. S3C), suggesting a compensatory expansion of remaining tissue-resident Vγ1^+^ T cells in the absence of Vγ7^+^ subset.

To address possible compensatory effects of tissue-resident Vγ1^+^ and Vγ7^+^ γδ T cells, we treated wild-type, Vγ7^-/-^ and Vγ7^+/-^ littermate control mice with depleting anti-Vγ1 antibody (clone 2.11) starting one week prior to AOM injection until the second DSS cycle. In contrast to untreated Vγ7^-/-^ mice, Vγ1–depleted Vγ7^-/-^ and Vγ7^+/-^ mice developed significantly more tumors, resulting in higher tumor load (Fig. 3A). Vγ1–depleted Vγ7^+/-^ and particularly Vγ1–depleted Vγ7^-/-^ mice display a significant reduction in γδ/αβ T cell ratio only in non-tumor areas, as well as in CD8aa^+^ γδ T cells (Fig. 3B to D). We did not observe differences in IFN-γ or IL-17 production among the groups but noticed an increase in PD-1–expressing γδ T cells in non-tumor areas, probably reflecting a relative increase in Vγ4^+^ or 6^+^ subsets (Fig. 3D and 3E). Overall, these results reinforce the idea that epithelium-resident Vγ1^+^ and Vγ7^+^ T cells play redundant anti-tumor activity.

**Figure 3.**
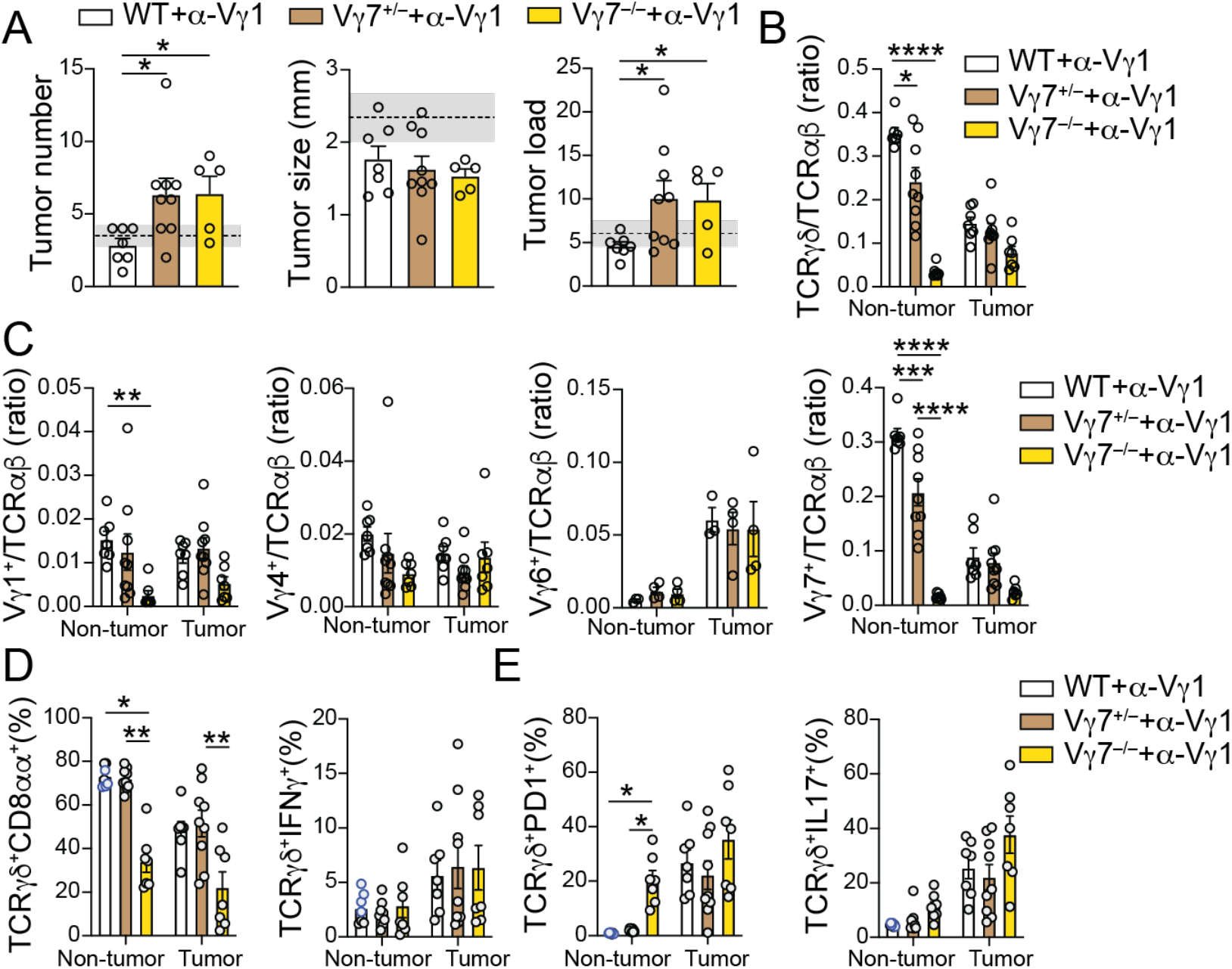
Tissue resident Vγ1^+^ and Vγ7^+^ γδ T cells synergize to control colon cancer development. (A-E) Wild-type B6 (WT), Vγ7-^/-^, and Vγ7^+/-^ littermate control mice treated with were subjected to AOM-DSS model and analyzed 12 weeks after initial AOM injection. All groups were treated with anti-Vγl depleting antibody (2.11) twice a week, starting one week before AOM administration until the second DSS cycle. (A) Tumor number, size and load. (B) Ratio of TCRγδ/αβ and (C) Vγl/TCRαβ, Vγ4/TCRαβ, Vγ6/TCRctβ, Vγ7/TCRαβ among CD45^+^ cells from colonic tissue. (D) Frequency of CD8α^+^ (left) and IFN-γ^+^ (right) and (E) PD-1^+^ (left) and IL-17^+^ (right) among TCRγδ^+^ cells. Data pooled from 2 experiments with 3-5 animals per group. * (p>0.05) ** (p>0.01) *** (p>0.001) symbols represent statical differences between groups.

While our observations point to an important anti-tumor function for epithelium–resident γδ T cells, previous studies have also described that tumor microenvironment–, or microbiota–influenced γδ T cell subsets could play an opposite role during tumor progression (*3, 24*). We therefore questioned whether the Vγ4^+^ and clonally expanded Vγ6^+^ PD-1^+^ IL-17 producing γδ T cells that accumulate among tumor infiltrating lymphocytes contribute to CRC progression. We conditionally deleted the main transcription factor linked to IL-17 production in T cells, Rorγt (*25*), using the same strategy described above. To specifically target γδ T cells that accumulate during CRC progression, we treated i*Trdc*^ΔRorc^ and littermate control mice with tamoxifen after the second DSS cycle (Fig. 4A). While no differences in tumor numbers or load were noted, late Rorγt deletion in γδ T cells results in smaller tumors (Fig. 4B). Accordingly, this strategy leads to altered Vγ-usage among tumor infiltrating γδ T cells: tamoxifen–treated i*Trdc*^*ΔRorc*^ mice display significantly decreased Vγ6^+^ and Vγ4^+^ populations when compared to littermate controls (Fig. 4C and fig. S4A). Tamoxifen-treated i*Trdc*^ΔRorc^ mice do not display changes in CD8αα-expressing γδ T cells (Fig. 4D). However, late Rorγt targeting results in reduced frequency of PD-1^+^ and IL-17–producing (71.2% and 39.08% suppression, respectively) tumor-infiltrating γδ T cells, affecting both Vγ6^+^ and Vγ4^+^ populations (4E and fig. S4B). No difference was noted in IL-17 production among CD4^+^ T cells (fig. S4C). These data indicate that Rorγt expression by γδ T cells is an important factor for the accumulation of IL-17–producing Vγ4^+^ and Vγ6^+^ TCRγδ^+^ cells and contributes to tumor growth.

**Figure 4.**
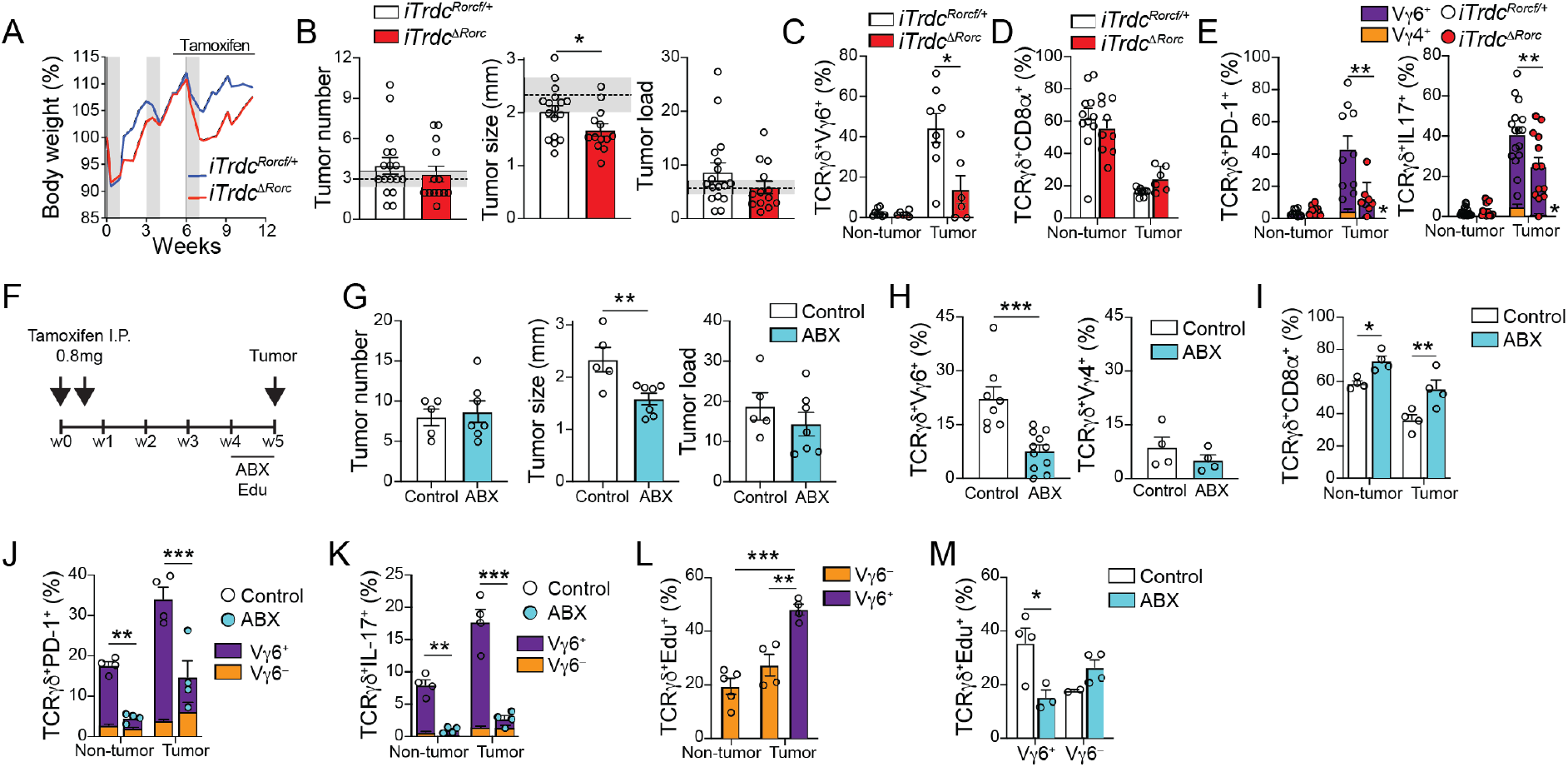
Loss-of-function by tumor-infiltrating IL-17^+^ γδ T cells results in smaller tumors. **(A-D)** *iTrdc*^Δ*Rorc*^ and littermate control (i*TrdC*^Δ*Rorcfl*+^) mice were subjected to AOM-DSS model, treated with tamoxifen for the last 6 weeks of the experiment and analyzed 12 weeks after initial AOM injection. **(A)** Percentage of body weight changes during AOM-DSS treatment. Gray bars represent DSS treatment. (B) Tumor number, size and load. Shaded area bounded by dashed lines indicates mean ± SEM of all control C57BL6/J mice analyzed in fig. S2B (AOM-DSS model). **(C-E)** Flow cytometry analysis of γδ T cells from tumor or non-tumor colonic tissue. **(C)** Frequency of Vγ6^+^ among TCRγδ^+^ cells. **(D)** Frequency of CD8α^+^ and **(E)** PD-1^+^ (left), and IL-17^+^ (right) cells among TCRγδ^+^ cells. Vγ6^+^ (purple) vs Vγ6 (orange) contribution to PD-1^+^ and IL-17–producing γδ T cells is also shown. **(F-M)** *iCdx2^ΔAPC^* mice were treated with 2 i.p. injections of O.8mg tamoxifen and analyzed 5 weeks after. Mice were treated with antibiotic mix (ABX) and EdU in the drinking water or only EdU (control) for the last week of the experiment. (F) Protocol. **(G)** Tumor number, size and load. **(H-M)** Flow cytometry analysis of γδ T cells from tumor or non-tumor colonic tissue. **(H)** Frequency of Vγ6^+^ (left) or Vγ4^+^ (right) among TCRγδ^+^ cells. **(I)** Frequency of CD8α÷ cells among TCRγδ^+^ cells. **(J)** Frequency of PD-1^+^ and (K) Il-17^+^ among TCRγδ^+^ cells. Vγ6^+^ (purple) vs Vγ6 (orange) contribution to PD-1^+^ and IL-17–producing γδ T cells is also shown. (L) Frequency of EdU incorporation by Vγ6 or Vγ6^+^ TCRγδ^+^ cells in tumor-containing colonic tissue isolated from control *iCdx2^ΔAPC^* mice. (M) Frequency of EdU incorporation by Vγ6^+^ and Vγ4^+^ TCRγδ^+^ cells in tumor-containing colonic tissue. i*Trdc*^*ΔRorc*^ tumor data are pooled from 3 experiments with 3-6 animals per group. Data from *iCdx2^ΔAFC^* are representative from 2 independent experiments with 3-4 animals per group. *P < 0.05, **P < 0.01, ***P < 0.001. **(C-E and I-M)** One-way ANOVA with Dunnett’s multiple comparison test; **(B, G, H)** two-tailed t-test. Error bars indicate SEM.

Changes in resident bacterial communities have been associated to tumor burden in CRC models (*9*) and were shown to impact pro-tumorigenic T cells, including Rorγt^+^ IL-17–producing Vγ6^+^ γδ T cells, in both lung and ovarian cancer models (*26, 27*). Due to complex changes in gut microbiota composition after cycles of DSS treatment (*27*), we focused on the impact of microbiota changes and antibiotic treatment in the APC loss model. 16S ribosomal RNA sequencing from feces of mice subjected to the APC loss model revealed a sharp decrease in microbial diversity as well as broad changes in bacterial composition that can be detected as early as 2 weeks after tamoxifen administration and continue in the following timepoints (fig. S4D and 4E). Consistent with a role of gut microbiota in promoting tumor growth (*9*), subjecting tamoxifen-treated *iCdx2^ΔAPC^* mice to a broad-spectrum antibiotic cocktail (ABX; Ampicillin, Vancomycin, Metronidazole and Neomycin) during the last week of the CRC development results in smaller tumors while not impacting tumor number or load, resembling changes observed upon late Rorγt targeting in the CAC model (Fig. 4F and 4G). Although microbiota depletion in tamoxifen-treated i*Cdx2*^ΔAPC^ mice leads to decreased intra-tumoral Vγ6^+^ γδ T cells, it does not affect Vγ4^+^ cells (Fig. 4H). CD8aa^+^ γδ T cells, primarily represented by Vγ1^+^ and Vγ7^+^ cells, display increased frequency in both non-tumor and tumor sites in mice receiving ABX treatment (Fig. 4I). Moreover, ABX treatment results in reduced frequency of IL-17– and PD-1–expressing γδ T cells (both over 75% suppression), primarily within the Vγ6^+^ subset (Fig. 4J and 4K), even though Vγ6^-^ γδ T cells (mostly comprised of Vγ4^+^) also contribute to PD-1 and IL-17 expression in tumor sites (see fig. S1H).

To address whether Vγ6^+^ cells accumulate during CRC progression due to increased recruitment or *in situ* proliferation, we performed *in vivo* EdU labelling. We observe higher proliferation rates in Vγ6^+^ *versus* Vγ6^-^ γδ T cells, which are reduced (56.7%) upon ABX treatment; Vγ4^+^ proliferation does not appear to be impacted by ABX treatment (Fig. 4L and 4M). These observations point to distinct susceptibility to microbiota manipulations by two main subsets of tumor-accumulating γδ T cells: Vγ6^+^, but not Vγ4^+^ cells, proliferate in response to microbiota and depend on microbiota signals to sustain PD-1 and IL-17 expression, as well as to boost tumor growth.

Our findings so far indicate that IL-17–producing, or PD-1–expressing tumor–infiltrating γδ T cells are composed ~ 75-80% of microbiota-dependent Vγ6^+^, and ~20-25% of Vγ4^+^ cells. To directly assess the roles of Vγ4^+^ and Vγ6^+^ γδ T cells in CRC progression, we generated Vγ4^-/-^ and Vγ6^-/-^ mice by CRISPR targeting of *Trgv4* and *Trgv6* genes, respectively. Fully backcrossed Vγ4^-/-^ mice were subjected to AOM-DSS and show no differences in tumor number, tumor size or tumor-infiltrating γδ T cells (aside from Vγ4) when compared to heterozygous littermate controls (fig. S5A-G). The ratio of γδ/αβ T cells among of tumorinfiltrating lymphocytes in Vγ4^-/-^ mice remains the same as the ratio observed in Vγ4^+/-^ controls (fig. S5B), suggesting a compensatory expansion of remaining γδ T cells, including Vγ6^+^ T cells, in the absence of Vγ4^+^ subset.

We next analyzed backcrossed naïve Vγ6^-/-^ mice. Like Vγ4^-/-^ mice, Vγ6^-/-^ mice subjected to AOM-DSS display similar CRC development, progression (Fig. 5A) and parameters of tumor-infiltrating γδ T cells to Vγ6^+/-^ littermate controls (Fig. 5B to F). The absence of an otherwise large intra-tumor Vγ6^+^ population did not result in altered γδ/αβ T cell ratio (Fig. 5B), explained by a compensatory increase in other γδ T cells, particularly Vγ4^+^ cells (Fig. 5C). We did not observe differences in tumor–infiltrating CD4^+^ T cells between Vγ6^-/-^ and Vγ6^+/-^ mice (fig. S5H). The overall similar tumor development and composition of tumorinfiltrating γδ T cells in Vγ4^-/-^ and Vγ6^-/-^ mice raise the possibility that tumor-infiltrating, PD-1^+^ IL-17–producing Vγ4^+^ and Vγ6^+^ populations play redundant roles in promoting CRC growth.

**Figure 5.**
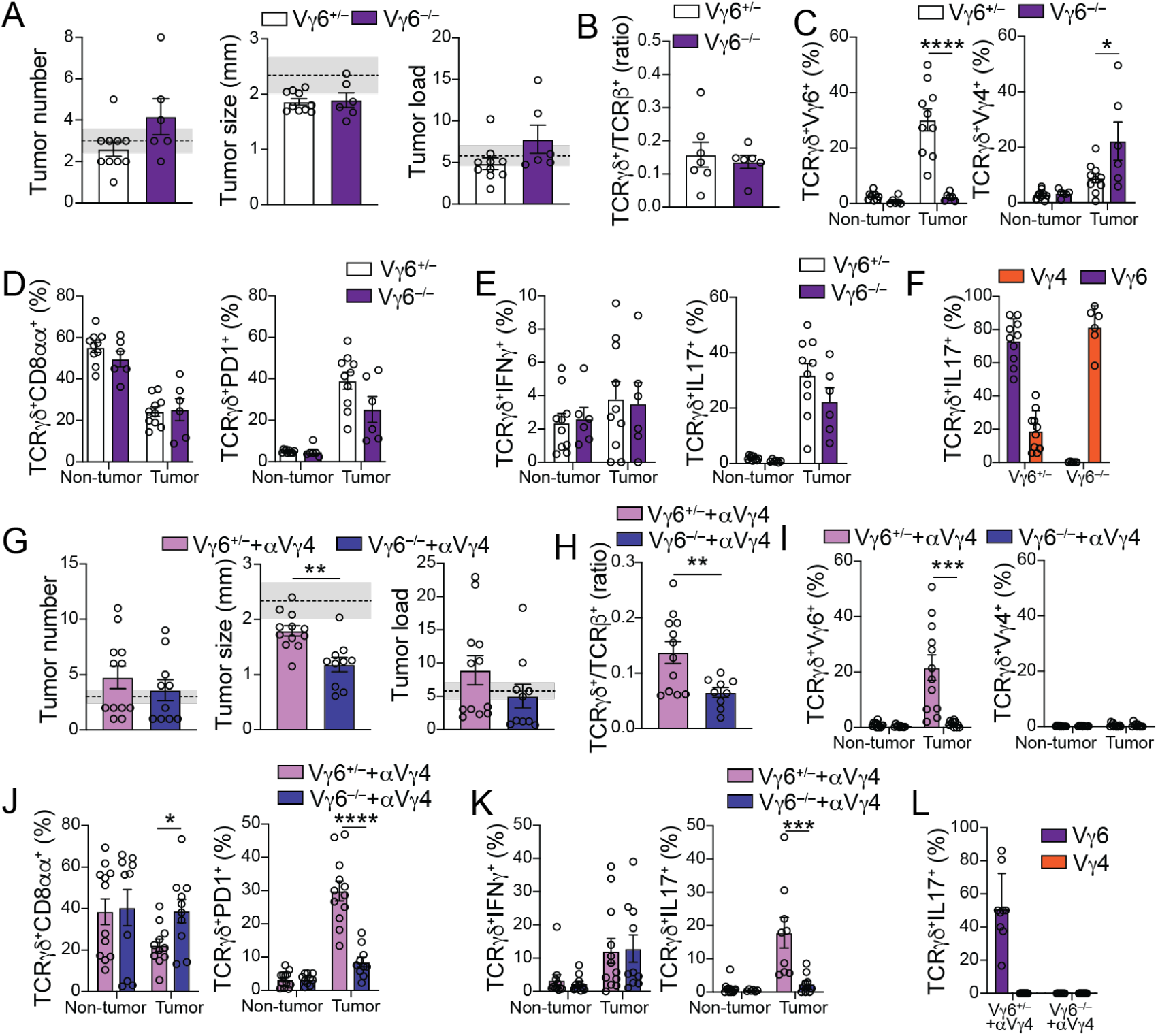
Redundant tumor-infiltrating IL-17–producing Vγ6^+^ and Vγ4^+^ γδ cells promote tumor growth. **(A-L)** Female Vγ6 and Vγ6^1^ littermate control mice were subjected to the AOM-DSS model and analyzed 12 weeks after initial AOM injection. In panels G to L, mice received injections of α-Vγ4 depleting antibody (UC3-10A6) twice a week starting one week after the 2nd DSS cycle (last six weeks of experiment). **(A, G)** Tumor number, size and load. Shaded area bounded by dashed lines indicates mean ± SEM of all control C57BL6/J mice analyzed in fig. S2B (AOM+DSS model). **(B-F; H-L)** Flow cytometry analysis of γδ T cells from tumor or non-tumor colonic tissue. **(B, H)** TCRγδ/αβ ratio among CD45^+^ cells from colonic tumor tissue. **(C, I)** Frequency of Vγ6^+^ (left) and Vγ4^+^ (right) and **(D, J)** CD8α^+^ (left) and PD-1^+^ (right) among TCRγδ^+^ cells from tumor or non-tumor colonic tissue. **(E, K)** Frequency of IFN-γ^+^ (left) and IL-17^+^ (right) among TCRγδ^+^ cells from tumor or non-tumor colonic tissue. **(F, L)** Frequency of Vγ6^+^ and Vγ4^+^ among IL-17–producing TCRγδ^+^ T cells in tumor-containing colonic tissue. Vγ6^-/-^ and anti-Vγ4-treated Vγ6 data are pooled from 2 experiments with 3-6 animals per group. *P < 0.05, **P < 0.01, *** P < 0.001. **(C-F; I-K)** One-way ANOVA with Dunnett’s multiple comparison test; **(A, B, G, H)** two-tailed t-test. Error bars indicate SEM.

To address possible compensatory, and redundant, roles between tumor–infiltrating Vγ4^+^ and Vγ6^+^ γδ T cells, we treated Vγ6^-/-^ and Vγ6^+/-^ littermate control mice with depleting anti-Vγ4 antibody (UC3-10A6) starting after the second DSS cycle until analysis. In contrast to untreated Vγ6^-/-^ mice, Vγ4–depleted Vγ6^-/-^ mice developed significantly smaller tumors than Vγ4-depleted Vγ6^-/+^ littermate control mice, while no significant changes in tumor numbers or load were noted (Fig. 5G). In contrast to previous strategies, Vγ4–depleted Vγ6^-/-^ mice display a roughly 50% reduction in tumor infiltrating γδ/αβ T cell ratio (Fig. 5H). Vγ4–depleted Vγ6^-/-^ mice also display enhanced intra-tumoral CD8aa^+^ γδ T cells when compared to Vγ4-depleted Vγ6^+/-^mice (Fig. 5J), similar to what was observed in ABX-treated mice in the APC loss model. Additionally, and consistent with a functional redundancy between Vγ4^+^ and Vγ6^+^ γδ T cells, Vγ4–depleted Vγ6^-/-^ mice show about 80% reduction in the frequency of PD-1^+^ and IL-17–producing γδ T cells within the tumor (Fig. 5J to L). No changes were observed in IFN-γ production, or in IL-17 production by CD4^+^ T cells, when compared to Vγ4–depleted Vγ6^+/-^ mice (fig. S5I). Hence, in sharp contrast to an anti-tumor role by epithelium-resident subsets, dominated by Vγ1^+^ and Vγ7^+/-^ γδ T cells, these data suggest redundant roles for tumor-infiltrating Vγ4^+^ and Vγ6^+^ γδ T cells in promoting CRC progression.

Tumor–infiltrating lymphocytes are essential components of anti-tumor responses and represent major targets for immunotherapies (*28–31*). Here we observed that γδ T cell subsets play contrasting roles in CRC: epithelial surveillance by steady-state IFN-γ–producing cytotoxic γδ populations help prevent tumor initiation, while accumulating intra-tumor γδ T cells support tumor progression. Specialized “dendritic epidermal T cells” (DETC) γδ T cells in the skin, mostly comprised of Vγ5^+^ cells have been shown to suppress tumor development via an NKG2D–dependent cytotoxic mechanism (*2, 32, 33*), while dermal γδ T cells, pre-committed to IL-17 production and expressing Vγ4 or Vγ6, were shown to promote tumor growth (*31*). While these observations parallel our findings, whether such anatomical segregation can also be observed in the intestinal epithelium *versus* lamina propria in specific conditions (*16*), remain to be determined. Our observations in the intestine indicate that γδ IELs in pre- or non-tumor areas harbor a diverse TCR repertoire while sharing primarily Vγ1 or 7 segments, particularly in the APC loss model. The accumulation of Vγ7^+^ IELs was previously linked to binding of the germline Vγ7 chain to Btnl proteins; a tissue-specific selection not associated with TCRγδ CDR3 region (*16*). Analogous to changes we observed in γδ T cells during CRC progression, previous studies described an irreversible expansion and repertoire reshaping of γδ T cells in chronically inflamed conditions such as in Celiac patients (*33*). Another important parallel with this study is their observation that γδ T cells from patients with active disease acquire a pro-inflammatory cytokine profile in contrast to γδ T cells isolated from patients in remission, which are primarily cytotoxic (*33*). Additionally, a recent study uncovered a metabolism-driven dichotomy in γδ T cell function: IL-17–producing γδ T cells are found to be dependent on oxidative phosphorylation, thriving in lipid-rich environments such as tumors; IFN-γ–producing γδ T cells required glycolysis for their energy expenditure (*23*). Conversely, previous observations suggested a role for clonally–restricted IL-17–producing T cells, including γδ cells, in tumor-progression (*9, 26, 27*). These observations are in line with our observations that, like their response to invading bacteria (*3*), IFN-γ–producing γδ T cells, particularly the Vγ7^+^ subset, display anti-tumor activity dependent on Glut1 expression, while IL-17–producing γδ T cells, particularly the Vγ6Vδ4 clone, were highly expanded in tumor areas. Our results, and previous literature (*9, 26, 31, 34–36*), point to a tumor-progression role for intra-tumoral IL-17. Whether downstream mechanisms induced by IL-17, such as neutrophil activity, or whether specific localization of these cells are necessary for tumor growth remain to be defined. Because IL-17 production by γδ T cells in the gut has also been linked to tissue repair (*37*), it remains possible that in addition to IL-17, other factors or molecules expressed by Vγ4^+^ or Vγ6^+^ cells aid tumor growth in the balance with tissue repair mechanisms. It would be relevant to define whether high PD-1 expression, specific localization within the tumor, or other factors, distinguish a tumor progression role for γδ T cells *versus* other IL-17–secreting cells, such as Th17 cells. Additionally, future studies are required to define the mechanisms by which the tumor microenvironment mediates the accumulation of microbiota–dependent Vγ6^+^ and microbiota–independent Vγ4^+^ subsets that aid tumor growth; it is tempting to speculate that analogous mechanisms to the Btnl–dependent Vγ7 selection (*15, 16, 22, 38*), are utilized. The results presented here caution against broad targeting of γδ T cells in future immune-therapy strategies, but they open possibilities of specific targeting of γδ T subsets based on Vγ-usage, pending confirmation of whether human CRC (*30*) parallels γδ T cell dynamics observed here.

## Acknowledgements

We thank all Mucida Lab members and Rockefeller University employees for their continuous assistance; A. Rogoz and S. Gonzalez for the maintenance of mice. We thank Rebecca O’Brien for the anti-Vγ7 hybridoma, and M. Constantinides & Y. Belkaid for anti-Vγ6 hybridoma. We also thank Victora and Lafaille labs for fruitful discussions. This work was supported by NIH R01CA218133 (S.G.), R01DK093674, R01DK113375, Food Allergy FARE/FASI Consortium, Mathers Foundation and Pershing Square Sohn Cancer Research Alliance (D.M.).

## Author Contributions

B.S.R. conceived, initiated, designed, performed, and analyzed experiments, and wrote the manuscript. P.D., A.L. and A. B. designed and performed experiments. I.Z.K., O.E., C.Z. and A. R. performed experiments. J.B. generated CRISPR mutant mice. T.B.R.C performed RNAseq and scTCRseq analysis. S.G. helped initiating, designing, and analyzing experiments. D.M. conceived, initiated, designed, supervised the research, and wrote the manuscript. All authors revised and edited the manuscript and figures.

## Supplementary Materials

### Methods

#### Mice

C57BL/6J control (000664), *Trdc^GFP^* (016941), *Trdc*^-/-^ (002120), *Trdc^CreER^* (031679), *Tbx21^fl/fl^* (022741), *Scl2a1^fl/fl^* (031871), *Rorc^fl/fl^* (008771), *Cdx2^CreER^* (022390), Rosa26^dTomato^ (007914) were purchased from Jackson Laboratories and kept in our facility. APC^*fl/fl*^ mice were provided by S. Grivennikov (C. Sinai) (*9, 12, 13*). E8_I_^CRE^ mice were provided by Ichicro Taniuchi (RIKEN, Japan). Vγ4^-/-^, Vγ6^-/-^, and Vγ7^-/-^ mice were developed by CRISPR-Cas9 gene editing targeting *Trgv4, Trgv6,* and *Trgv7* genes (*see below*). Heterozygous or wild-type homozygous littermates were used as controls. Mice were bred within our facility to obtain strains described and were 7-12 weeks of age for all experiments. To minimize microbiome fluctuations in the AOM-DSS model, genetically modified and wild-type control mice were co-housed from birth. Male and female mice were used in all experiments, in similar distribution, unless otherwise stated in the figure legends. Estrous cycle was not controlled for female mice. Animal care and experimentation were consistent with NIH guidelines and approved by the Institutional Animal Care and Use Committee (IACUC) at The Rockefeller University. All animals were kept in specific pathogen free (SPF) conditions.

#### Generation of Vγ-specific knockout strains by CRISPR-Cas9 gene editing

gRNA targeting specific *Trvg* genes were designed and injected into one cell stage fertilized C57BL/6J embryos together with Cas9 protein. Embryos were implanted into pseudo-pregnant foster Swiss mice and mutant offspring were selected by specific PCR genotyping; mutations were determined by sanger DNA sequencing. Mutant mice were backcrossed to wild-type C57BL/6J animals for 5-10 generations. For *Trvg4* gene targeting sgRNA sequence GUGUAACCAUACACUGGUACCGG was used. For *Trvg6* gene targeting, sgRNA AGCCCGAUGCAUACAUACACUGG and for *Trvg7* gene targeting sgRNA sequence ACUGGUACCGAUUCCAGAAAGGG was used. Mice were kept in our facility under the Rockefeller University IACUC protocol.

### Colorectal cancer (CRC) models

#### AOM-DSS model

Mice were injected with 12.5mg/kg of Azoxymethane (AOM) 1 day prior to 3 consecutive oral exposures of dextran sodium sulfate (DSS, 2% in drinking water) for 1 week with 2 weeks interval between exposures. Animals were sacrificed 12 weeks after initial AOM injection. Tumors were counted and measured at the time of experiment termination.

#### APC model

*Cdx2^CreER^*:APC^fl/fl^ (*iCdx2*^ΔAPC^) mice were injected twice with 0.8 mg of tamoxifen (Sigma) 4mg/ml in corn oil (Sigma), two days apart. Four weeks after the first injection, mice were sacrificed, and tumors were measured and counted. Animals were monitored for weight loss and sickness behavior and the experiment was terminated when weight loss was greater than 20% of initial weight.

#### Tamoxifen treatment regimen in AOM-DSS model

Tamoxifen (Sigma) was administrated by intraperitoneal injections, twice a week, of 0.8mg of tamoxifen diluted 4 mg/mL in corn oil (Sigma). For early gene depletion, tamoxifen injections started 1 week before AOM administration and continued until week 3 of AOM-DSS CRC model. For late gene targeting, tamoxifen injections started 1 week after second round of DSS and continued until the week 12 of the AOM-DSS CRC model.

#### Antibiotic and EdU administration

*iCdx2^ΔAPC^* mice were treated for one week with broad spectrum antibiotics consisting of 1mg/ml ampicillin, 1mg/ml neomycin, 0.5mg/ml vancomycin, 0.5mg/ml metronidazole and 10g/ml sucralose in the drinking water during the final week of the experiment. EdU was also added to the antibiotic containing water at 1mg/ml. As a control, 10mg/ml sucralose solution containing 1mg/ml EdU was used. EdU detection was performed using the Click-iT™ Plus EdU Flow Cytometery Assay kit (Thermo Fisher Scientific, C10632), according to manufacturer’s instructions.

#### *In vivo* antibody administration

For depletion of Vγ4^+^ T cells, 200μg of anti-Vγ4 antibody UC3-10A6 (Bio X Cell) was injected intra peritoneally twice a week for the final 6 weeks of AOM-DSS model. For antibody depletion of Vγ1^+^ T cells, 200μg of anti-Vγ1 antibody 2.11 (Bio X Cell) was injected as described for UC3-10A6.

#### Isolation of intestinal and tumor infiltrating cells

Intraepithelial lymphocytes were isolated as previously described (*39*). Briefly, large intestines were harvested and washed in PBS and 1mM dithiothreitol (DTT) followed by 30 mM EDTA. Intraepithelial cells were recovered from the supernatant of DTT and EDTA washes. Tumor infiltrating cells were obtained after collagenase digestion of the tissue. Dissected tumor was minced and incubated with 50mg/ml of collagenase VIII, 200mg/ml of DNAse in RPMI for 30 minutes. Mononuclear cells were isolated by gradient centrifugation using Percoll. Single-cell suspensions were then stained with fluorescently labeled antibodies for 25min at 4°C prior to flow cytometry analysis and/or cell sorting.

#### Flow cytometry

Cells were stained for anti-CD45 (BD 561487 clone 30-F11), anti-TCRβ (BioLegend 109220 clone H57-597) anti-TCRγδ (eBioscience 11-5711-85 clone GL-3), anti-CD8α, (eBioscience 56-0081-82 clone 53-6.7), anti-CD4 (BioLegend 100552 clone RM4-5), anti-PD-1 (BioLegend 135219 clone 29F.1A12), anti-Vγ1 (BioLegend 141109 clone 2.11), anti-Vγ4 (eBioscience 25-5828-82 clone UC3-10A6), anti-Vγ6 (clone 17D1, kindly donated by Yasmin Belkaid, Ph.D.), anti-Vγ7 (clone GL1.7, kindly donated by Rebecca O’Brien, Ph.D.), anti-CD107 (eBioscience 46-1071-82 clone 1D4B), anti-IFN-γ (eBioscience 45-7311-82 clone XMG1.2), anti-IL-17a (BD 559502 clone TC11-18H10), anti-CD11b (eBioscience 47-0112-82 clone M1/70). For anti-Vγ6 staining, cells were first stained with anti-TCRγδ (clone GL-3) for 30 minutes prior anti-Vγ6 17D1 antibody staining. For cytokine profiling, tissue isolated cells were stimulated ex-vivo with 1mg/ml of phorbol myristate acetate (PMA) and 1mg/ml of Ionomycin in the presence of 10mg/ml of Brefeldin for 4 hours prior staining.

#### Bulk mRNA-Seq Library Preparation

Sorted cells (300-800 cells) were lysed in a guanidine thiocyanate buffer (TCL buffer, QIAGEN) supplemented with 1 % β-mercaptoethanol. RNA was isolated by solid-phase reversible immobilization bead cleanup using RNAClean XP beads (Agentcourt, A63987), reverse transcribed, and amplified as described (*40*). Uniquely barcoded libraries were prepared using Nextera XT kit (Illumina) following manufacturer’s instructions. Sequencing was performed on an Illumina NextSeq500 for a total yield of 400M reads.

#### Bulk RNA-Seq Analysis

Raw fastq files were processed by using the mouse transcriptome (gencode M23) with the kallisto (v0.46) software (*41*). Analysis of transcript quantification was performed at the gene level by using the sleuth (v0.30) package for R (*42*). Shortly, we modeled batch effect and our experimental design using the sleuth_fit function and detected differentially expressed genes between all groups by the likelihood ratio test (LRT). To spot statistically significantly enriched genes between group pairs, we used the wald-test function. Genes with false discovery rate less than 0.05 and 1 log2 fold-change were used in downstream analysis. The batch effect was removed from the expression matrix by using the *removebatcheffect* function available in the limma package (*43*). Gene set enrichment analysis (GSEA) was performed by using signature gene sets in gmt format and a pre-ranked gene list by log2 fold-change between two groups as input for the fgsea. Gene ontology (GO) analysis was executed by comparing all detected genes as our background and the lists of differentially expressed genes against the biological processes gene sets by using topGO (*44, 45*).

#### Single-Cell TCR Sequencing

Single cells were index-sorted using a FACS Aria into 96-well plates containing 5 μL of lysis buffer (TCL buffer, QIAGEN 1031576) supplemented with 1 % β-mercaptoethanol and frozen in −80°C prior to RT-PCR. RNA and RT-PCRs for TCRγ and TCRδ were prepared using specific primers (supplemental table 1). PCR products for TCRγ and TCRδ were multiplexed with barcodes and submitted for MiSeq sequencing (*46*) using True Seq Nano kit (Illumina). Fastq files were de-multiplexed and paired-end reads were assembled at their overlapping region using PANDASEQ (*47*) and FASTAX toolkit. De-multiplexed and collapsed reads were assigned to wells according to barcodes. Fasta files were aligned and analyzed on IMGT (http://imgt.org/HighV-QUEST) (*48*). Cells with identical TCRγ and TCRδ CDR3 nucleotide sequences were considered as the same clones. Clonality was assigned based on paired TCRγδ per mouse. Only in frame junction sequences of TCRγδ were included in the analysis.

#### FocusClear and cell localization on villus-crypt axis

Tissue clearing was performed according to FocusClear manufacturer’s (Celexplorer Labs. Co.) instructions. Briefly, E8_I_ (*Cd8a* enhancer I)^Cre^:*Rosa26*^fsf-dTomato^ (E8_I_^dTomato^):*Trdc*^GFP^ double-reporter mice were injected with Hoechst dye (blue) for visualization of epithelial cell nuclei. After 15-30 min, mice were sacrificed by cervical dislocation and segments of the small intestine were removed, washed (intact) and fixed in 4% PFA at gentle agitation for 2h at RT. After fixation, samples were washed in DPBS and placed in FocusClear solution for approximately 15 min. at room temperature. Once visual confirmation of clearance was obtained, samples were mounted in 3D printed slides with MountClear, sealed and imaged using an inverted LSM 880 NLO laser scanning confocal and multiphoton microscope (Zeiss). As this protocol maintains native GFP and dTomato, and introduced Hoechst fluorescence, no antibody staining was necessary. Cell X and Y coordinates were obtained using IMARIS (Bitplane) software, as described before (*3*). Multiple technical replicates per sample (segment) were obtained.

#### Statistical Analyses

Statistical analyses were carried out using GraphPad Prism v.9. Flow cytometry analyses were carried out using FlowJo software. Data in graphs show mean ± SEM and P values < 0.05 were considered significant. Statistical analyses were performed using ordinary one-way ANOVA test with Dunnett’s multiple comparison test, two-tail t-test or Chi-square test, as indicated in the figure legends. *P < 0.05, **P < 0.01, ***P < 0.001. GraphPadPrism v.9 was used for graphs and statistical analysis, and Adobe Illustrator 2020 was used to assemble and edit figures.

## Supplementary Figures

**Figure S1. Supporting data to Figure 1.**
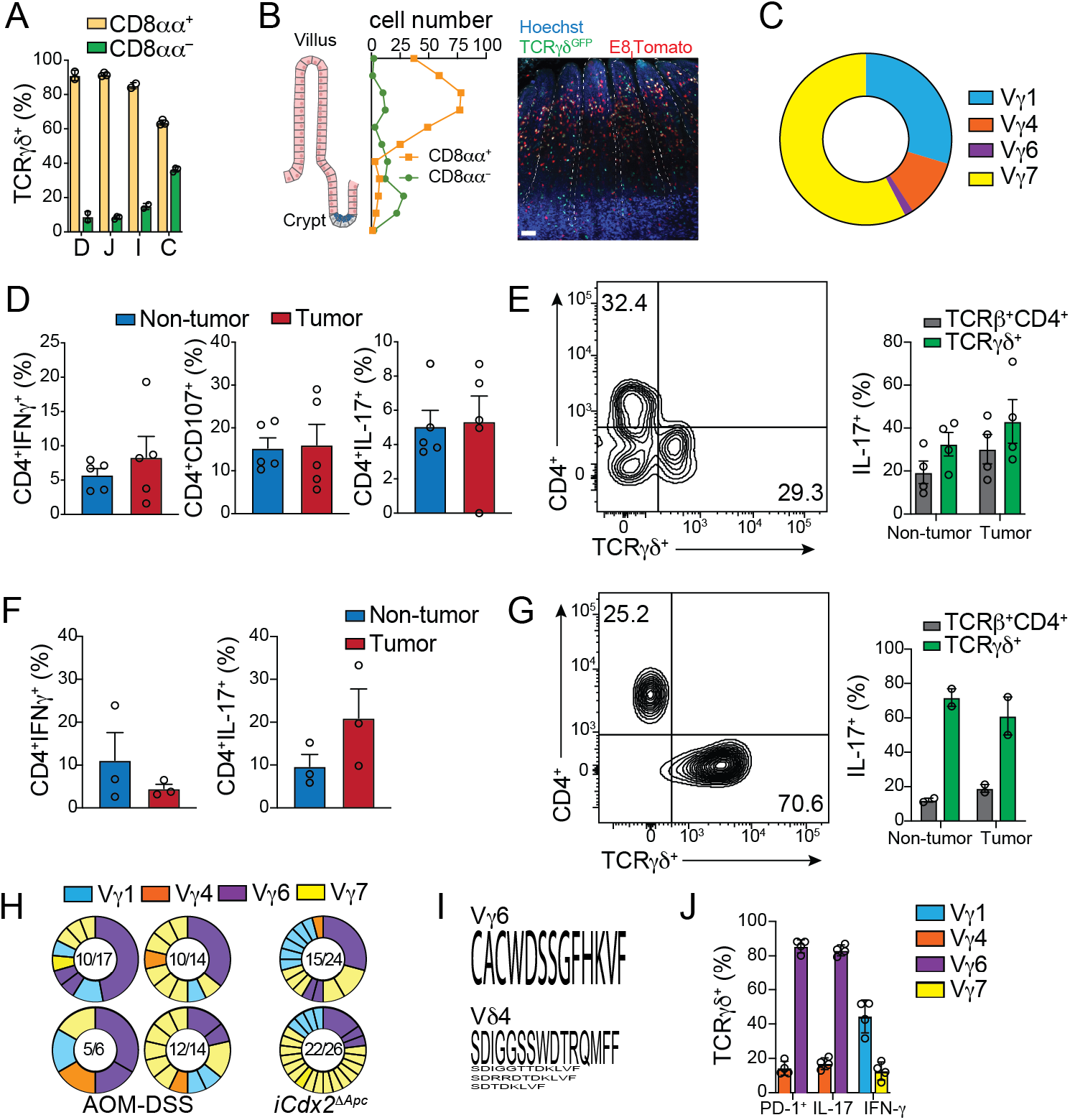
(A, C) Flow cytometry analysis of naïve C57BL/6 mice. (A) Frequency of CD8αα^+^ and CD8αα^-^ among TCRγδ^+^ cells from duodenum D, jejunum J, ileum I and colon C. (B) Whole-mount imaging of tissue-cleared ileum from E8_l_^Tomato^Trdc^GFP^ reporter mouse at steady-state. Image shows TCRγδ^+^CD8αα^+^ (Tomato^+^GFP^+^) and TCRγ-δ^+^CD8αα^-^ (Tomato-GFP^+^) cells along the villus-crypt axis. Scale bar= 4Oμm. (C) Frequency of Vγ usage by colonic TCRγδ^+^ cells at steady state. (D, E) Flow cytometry analysis of γδ T cells from tumor or non-tumor ∞lonic tissue of C57BL/6 mice subjected to AOM-DSS model. (D) Frequency of IFN-γ^+^ (left), CD107^+^ (center) and IL-17^+^ (right) among TCRαβ^+^CD4^+^ cells. (E) Representative dot-plot (left) and frequency of CD4^+^ and TCRγδ^+^ cells among IL-17^+^ cells. (F, G) Flow cytometry analysis of γδ T cells from tumor or non-tumor colonic tissue of tamoxifen-treated iCdx2Δ^APC^. (F) Frequency of IFN-γ^+^ (left) and IL-17^+^ (right) among TCRαβ^+^CD4^+^ cells from colonic tumor. (G) Representative dot-plot (left) and frequency (right) of CD4^+^ and TCRγδ^+^ cells among IL-17^+^ cells. (H, I) Single-cell TCR sequencing of γδ T cells from tumor or non-tumor colonic tissue of mice subjected to the AOM-DSS (left, center) and APC loss (right) models. (H) Numbers in the center of pie charts represent number of clones per total cells sequenced. Expanded clones are fused. Clones are colored based on Vγ usage. Purple clones represent expanded Vγ6Vδ4 cells. Each pie chart represents an individual mouse. (I) Amino acid sequence used by Vγ6Vδ4 expanded clones (purple). Font size represents frequency of usage. (J) Frequency of Vγ1^+^, Vγ4^+^, Vγ6^+^ and Vγ7^+^ among tumor-infiltrating γδ T cells expressing PD-1, IL-17 or IFN-γ (APC loss model). Data representative from 2 experiments with 3-5 animals per group. *P < 0.05, **P < 0.01, ***P < 0.001. (A, C, D, F) One-way ANOVA test with Dunnett’s multiple comparison test; (E, G, J) two-tailed t-test. Error bars indicate SEM.

**Figure S2. Supporting data to Figure 2.**
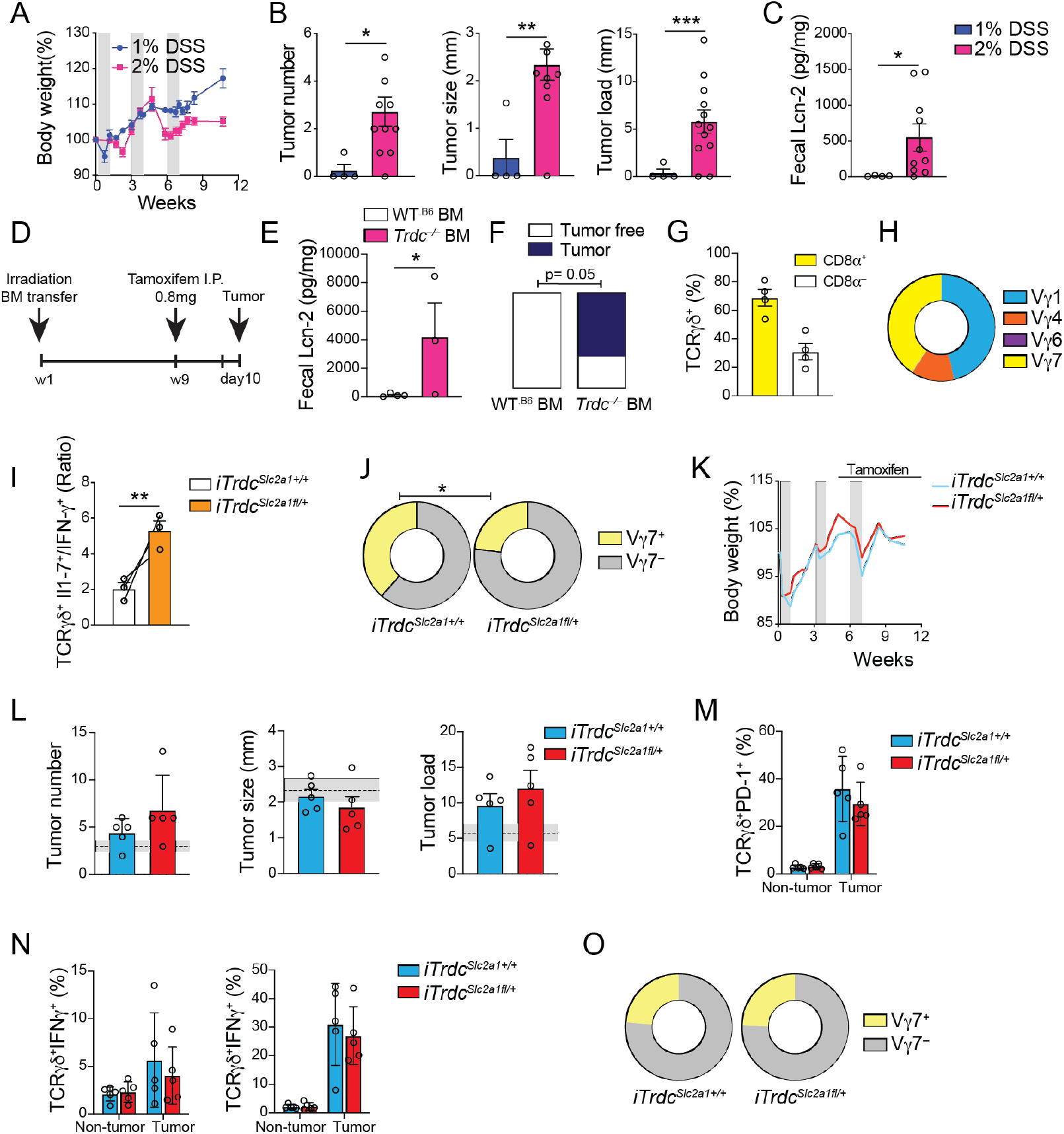
(A-C) C57BL/6 mice were injected with 12.5mg/kg of Azoxymethane (AOM) 2 days prior to three rounds of 1 week “on” and 2 weeks “off’ of 1% vs 2% Dextran sodium sulfate (DSS) in the drinking water. (A) Percentage of body weight changes during AOM-DSS treatment. Gray bars represent DSS treatment. (B) Tumor number, size and load. (C) Fecal lipocalin-2 quantification. (D-H) iCdx2Δ^APC^ mice were lethally irradiated (900 rad) and transferred with bone marrow from Trdc^-/-^ or control wild-type B6 animals. Eight weeks after bone marrow transfer, iCdx2Δ^APC^ recipient mice were injected with a single dose of 0.8 mg tamoxifen and analyzed 10 days later. (D) Protocol. (E) Fecal lipocalin-2 quantification. (F) Quantification of tumors in the colon. (G, H) Flow cytometry analysis of γδ T cells from non-tumor colonic tissue of control (WT.B6) bone marrow reconstituted iCdx2Δ^APC^. (G) Frequency of CD8α^+^ and CD8α^-^. (H) Vγ usage among TCRγδ^+^ cells. (I, J; K-O) iTrdc^Slc2a1fl/+^ and iTrdC^ΔScl2a1+/+^ littermate control mice were subjected to AOM-DSS model, treated with tamoxifen for the first 3 weeks (I, J) or the last 6 weeks (K-O) and analyzed 12 weeks after initial AOM injection. (I, J) Flow cytometry analysis of intra-tumor γδ T cells in colon. (I) Ratio of IL-17^+^ to IFN-γ^+^ cells among TCRγδ^+^ cells. (J) Frequency of Vγ usage by TCRγδ^+^ cells. (K) Percentage of body weight changes during AOM-DSS treatment. Gray bars represent DSS treatment. (L) Tumor number, size and load. Shaded area bounded by dashed lines indicates mean ± SEM of all ∞ntrol C57BL6/J mice analyzed in fig. S2B (AOM+DSS model). (M-O) Flow cytometry analysis ofγδ T cells from tumor or non-tumor colonic tissue. (M) Frequency of PD-1^+^ cells among TCRγδ” cells. (N) Frequency of IFN-γ^+^ (left) and IL-17^+^ (right) among TCRγδ^+^ cells. (O) Frequency of Vγ usage by tumor infiltrating TCRγδ^+^ cells. iCdx2Δ^APC^ bone marrow transfer and iTrdc^Slc2a1fl/+^ data representative from 2 experiments with 3-5 animals per group. 1% vs 2% DSS data are pooled from 2 experiments with 2-5 animals per group. *P < 0.05, **P <0<.01, ***P < 0.001. (M, N) One-way ANOVA test with Dunnett’s multiple comparison test; (B-E; G-J; O) Two-tailed t-test; (F) two-tailed Chi-test. Error bars indicate SEM.

**Figure S3. Supporting data to Figure 3.**
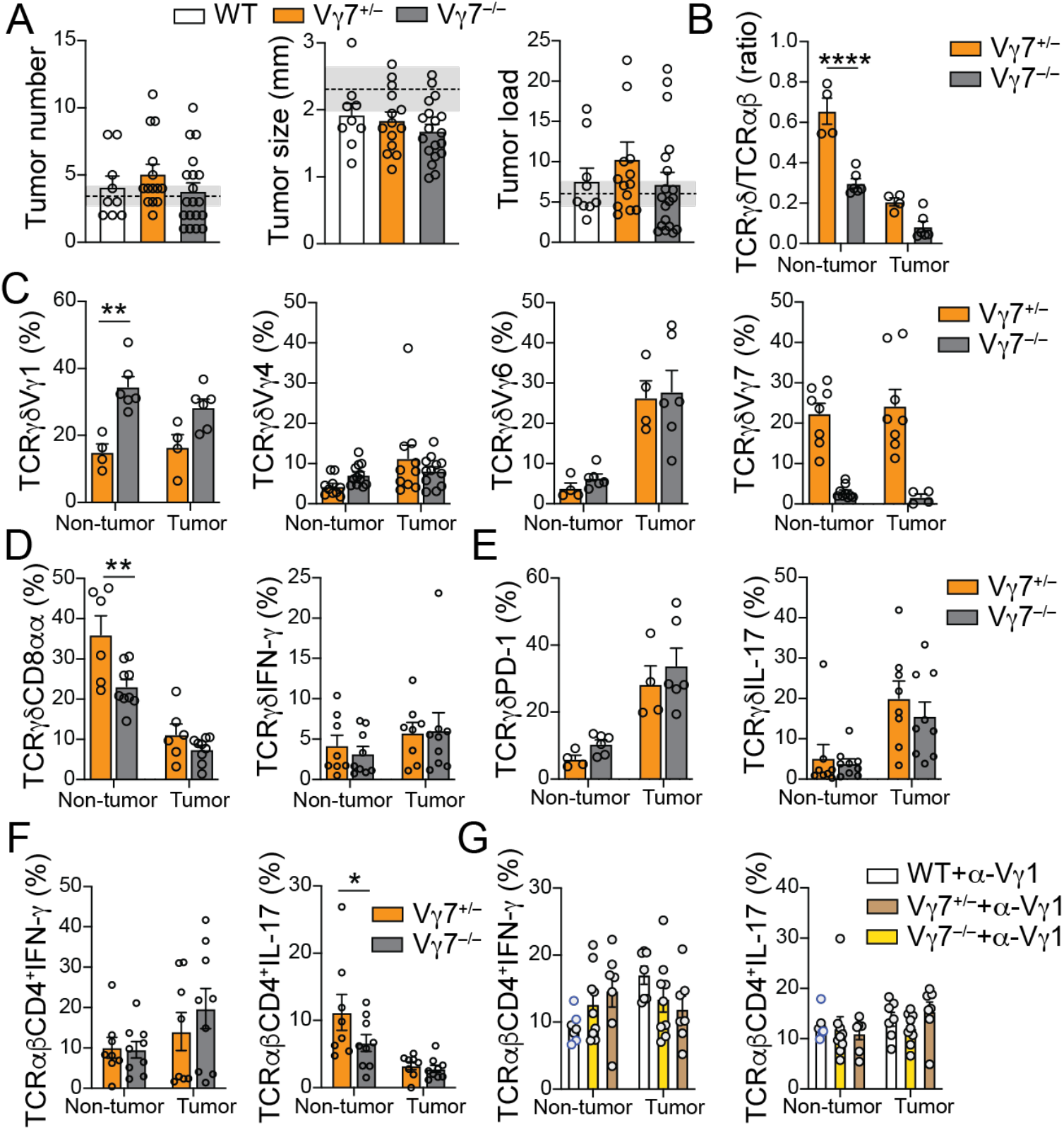
**(A-G)** Wild-type B6 (WT), Vγ7^-/-^, and Vγ7^+/-^ littermate control mice treated with were subjected to AOM-DSS model and analyzed 12 weeks after initial AOM injection. **(A)** Tumor number, size and load. **(B)** TCRγδ/αβ ratio and **(C)** frequency of Vγl, Vγ4, Vγ6, Vγ7 among TCRγδ^+^ cells from colonic tissue. (D) Frequency of CD8ct^+^ (left) and IFN-γ^+^ (right) and **(E)** PD-1^+^ (left) and IL-17* (right) among TCRγδ^+^ cells. **(F)** Frequency of IFN-γ* and IL-17* among CD4* T cells. (G) All groups were treated with anti-Vγl depleting antibody (2.11) twice a week, starting one week before AOM administration until the second DSS cycle. Frequency of IFN-γ* (left) and IL-17* (right) among CD4* T cells. Data pooled from 3 (A-F) and 2 (G) experiments with 3-5 animals per group. * (p>0.05) ** (p>0.01) *** (p>0.001) symbols represent statical differences between groups.

**Figure S4. Supporting data to Figure 4.**
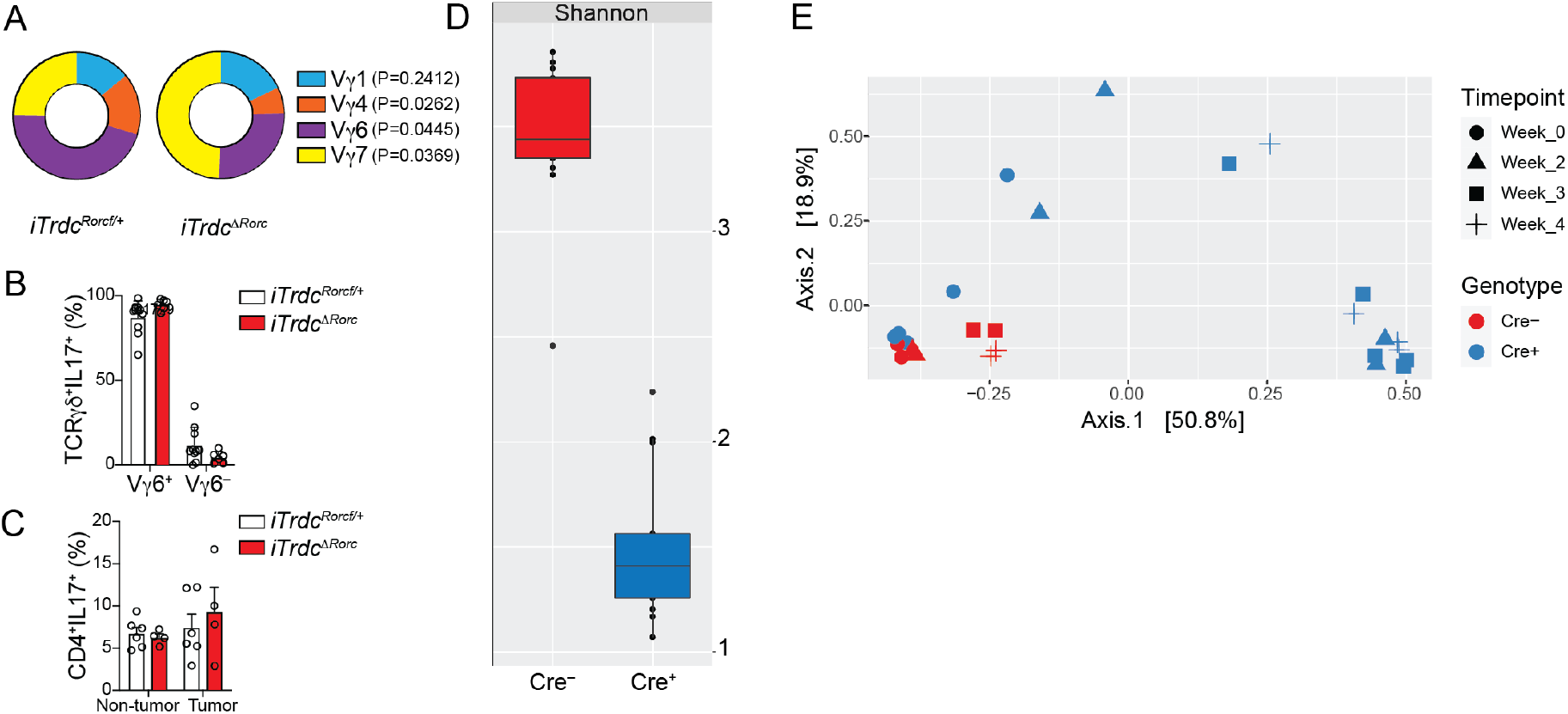
(A-C) iTrdc^ΔRorc^ and iTrdc^Rorcfl/+^ littermate ∞ntrol mice were subjected to AOM-DSS model, treated with tamoxifen for the last 6 weeks of the experiment and analyzed 12 weeks after initial AOM injection. Flow cytometry analysis of γδ T cells from tumor and non-tumor colonic tissue. (A) Frequency of Vγ usage among TCRγδ^+^ cells isolated from tumor colonic tissue. (B) Frequency ofVγ6^+^ vs Vγ6^-^ among IL-17–producing TCRγδ^+^ cells isolated from tumor colonic tissue. (C) Frequency of IL-17^+^ among TCRαβ^+^ CD4^+^ cells. (D, E) Bacterial 16S ribosomal RNA amplicon (16S rRNA) sequencing from fecal pellets of tamoxifen-treated iCdx2^ΔAPC^ and APC^f/f^ littermate ∞ntrol mice. (D) Shannon alpha diversity plot from iCdx2^ΔAPC^ (Cre^+^ blue) and APC^f/f^ control animals (Cre^-^ red), four weeks after tamoxifen administration. (E) Bi-dimensional principal coordinate analysis plot from iCdx2^ΔAPC^ (Cre^+^ blue) and Apc^f/f^ control animals (Cre^-^ red) before (week 0), 2, 3 and 4 weeks after tamoxifen administration. 16S rRNA sequencing data representative of 2 experiments with 2-4 animals per group. iCdx2^ΔAPC^ and APC^f/f^ control mice were co-housed. Fecal samples were collected from mice housed in different cages. (A) two-tailed t-test (P values are written in the figure); (B, C) One-way AN OVA test with Dunnett’s multiple comparison test. Error bars indicate SEM.

**Figure S5. Supporting data to Figure 5.**
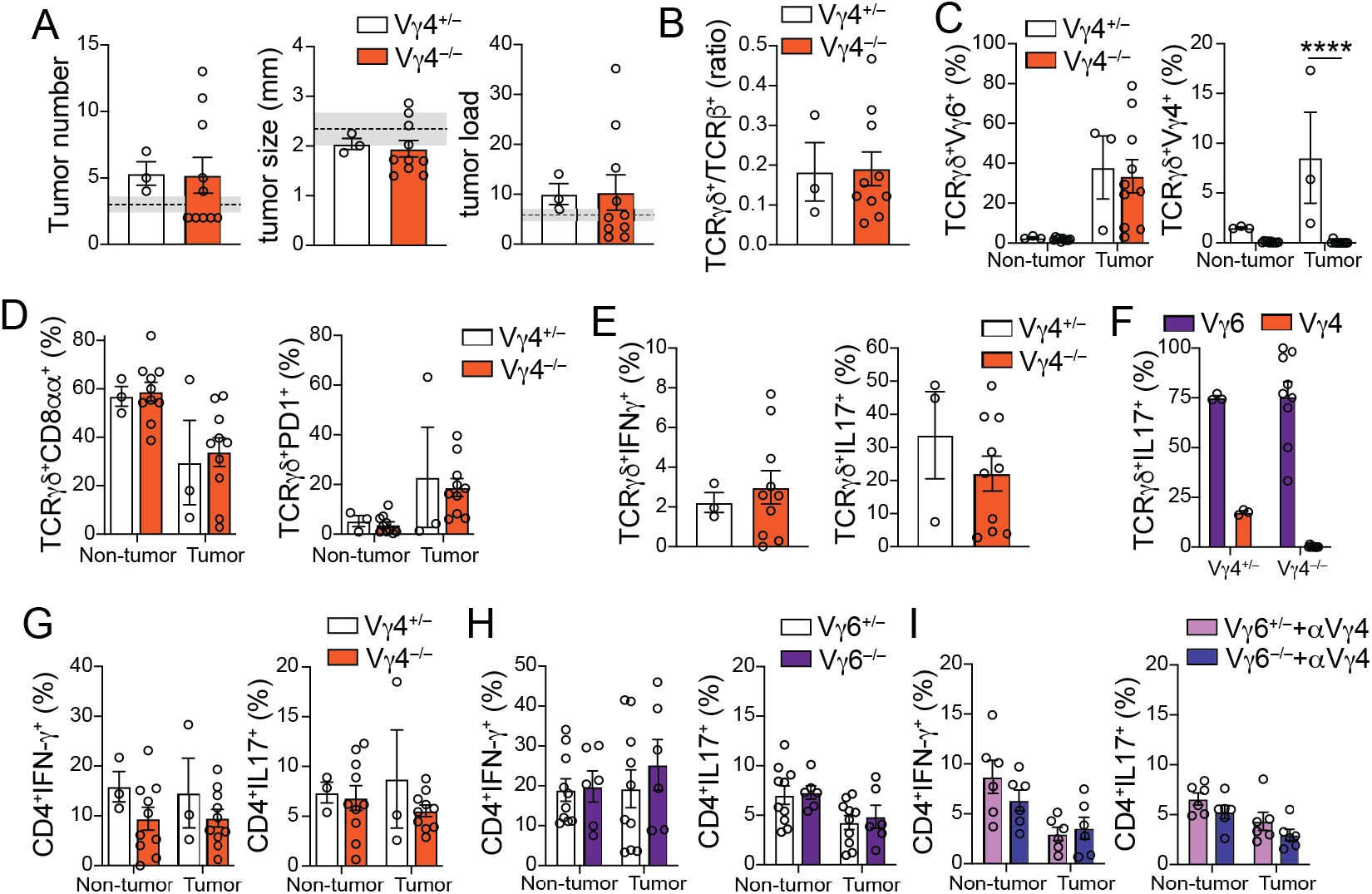
(A-G) Female Vγ4^-/-^ and Vγ4^+/-^ littermate control, (H, I) Vγ6^-/-^ and Vγ6^+/-^ littermate control mice were subjected to the AOM-DSS model and analyzed 12 weeks after initial AOM injection. In panel I, mice received injections of α-Vγ4 depleting antibody (UC3-10A6) antibody twice a week starting one week after the 2nd DSS cycle (last six weeks of experiment). (A) Tumor number, size and load. Shaded area bounded by dashed lines indicates mean ± SEM of all control C57BL6/J mice analyzed in fig. S2B (AOM+DSS model). (B-F) Flow cytometry analysis of γδ T cells from tumor or non-tumor ∞lonic tissue. (B) TCRγδ/αβ ratio within CD45^+^ cells from tumor colonic tissue. (C) Frequency of Vγ6^+^ (left) and Vγ4^+^ (right), and (D) CD8α^+^ (left) and PD-1 ^+^ (right) among TCRγδ^+^ cells. (E) Frequency of IFN-γ^+^ (left) and IL-17^+^ (right) among TCRγδ^+^ cells from tumor colonic tissue. (F) Frequency of Vγ6^+^ and Vγ4^+^ among IL-17–producing γδ T cells in tumor colonic tissue. (G-l) Frequency of IFN-γ^+^ (left) and IL-17^+^ (right) among TCRαβ^+^CD4^+^ cells from tumor or non-tumor colonic tissue of Vγ4^-/-^ (G), Vγ6^-/-^ (H) or Vγ4-depleted Vγ6^-/-^ (I) and littermate heterozygous control mice. Vγ4^-/-^ data pooled from 2 experiments with 3-5 animals per group. *P < 0.05, **P < 0.01, ***P < 0.001. (C, D; F-l) One-way ANOVA test with Dunnett’s multiple comparison test; (A, B, E) two-tailed t-test. Error bars indicate SEM.

